# Uplift and erosion of genomic islands with standing genetic variation

**DOI:** 10.1101/057513

**Authors:** Tyler D. Hether

## Abstract

Details of the processes that generate biological diversity have long been of interest to evolutionary biologists. A common theme in nature is diversification via divergent selection with gene flow. Empirical studies on this topic find variable genetic differentiation throughout the genome, that genetic differentiation is non-randomly distributed, and that loci of adaptive significance are often found clustered together within “genomic islands of divergence”. Theoretical models based on new mutations show how these genomic islands can arise and grow as a result of a complex interaction of various evolutionary and genic processes. In the current study, I ask if such genomic islands can alternatively arise from divergent selection from standing genetic variation and I tested this using a simple two locus model of selection. There are numerous ways in which standing genetic variation can be partitioned (e.g., between alleles, between loci, and between populations) and I tested which of these scenarios can give rise to an island pattern compared to no genomic differentiation or complete genomic differentiation. I found that divergent selection, even without reciprocal gene exchange between populations, following a bout of admixture can relatively quickly produce an island pattern. Moreover, I found two pathways in which islands can form from divergence from standing variation: 1) through the build up of islands and 2) through the breakdown of larger, genome-wide differentiation. Lastly, similar to new mutation theory, I found that the frequency of recombination is an important determinant of island formation from standing genetic variation such that mating behavior of a species (e.g., facultative or obligate sexual) can impact the likelihood of island formation.

## 1. Introduction

It is increasingly evident that phenotypic and taxonomic diversity arises despite ongoing gene flow between populations or incipient species [15]. Predicting the genomic response to divergence with gene flow (DGF) in nature is difficult, however, because several interacting evolutionary and genetic factors can occur simultaneously. Moreover, some of these factors can themselves have multiple levels of interaction. For example, divergent selection contributes to genetic divergence both *directly* by its effect on actual selected loci and *indirectly* by ‘divergent hitchhiking’ (DH) of nearby neutral loci [16].

The metaphor of “genomic islands of divergence” has been used recently to integrate the dynamics of migration and divergent selection affecting selected loci and recombination and selection affecting the degree of genetic hitchhiking [14, 10]. Here, inter-population gene flow homogenizes the neutrally evolving “sea floor” whereas DH creates genomic isolation, reducing the effective migration rate at selected loci as well as loci in tight physical linkage with these selected loci [16]. Such reduction in effective migration owing to DH can further diverge weakly selected, *de novo* mutations at nearby loci [18] that would otherwise be trumped by migration experienced at the sea floor. Thus, over time these divergent islands are hypothesized to grow (widen) with the inverse of the product of migration and recombination whereas height (extent of differentiation) is expected to be proportional to strength of divergent selection.

Mathematical models of GI formation has almost exclusively focused on divergent selection based on new mutations even though many research programs find adaptation from standing genetic variation [SGV; 13, 3, 2, 8, 7, 9]. Adaptation from SGV can lead to faster evolution, fixation of more small effect alleles, and an increase frequency of beneficial recessive alleles [11] relative to adaptation from new mutations [6, 1]. With regards to GI architecture, however, less is known about the role of SGV in part because such variation can be partitioned several different ways both within and between populations. For example, two populations might be fixed for alternate alleles at all polymorphic loci such that each population is in linkage equilibrium but there is a high degree of cross-population linkage disequilibrium (X-LD) between loci. In such a case all the SGV is partitioned between populations. In other cases, a varying level of polymorphism can occur within one or more populations at one or more loci. It is reasonable to suspect that varying how SGV is partition would likely affect the overall magnitude and localization of genetic differentiation nearby loci under divergent selection.

The arrangement of genetic differentiation that occurs across the genome varies widely in the literature [10] which makes drawing conclusions on the nature of genomic differentiation difficult. It has been postulated that islands form by DH, with growth of such chromosomal regions possible by further divergent selection occurring at loci that are themselves linked to an already established divergently selected locus [10, 18]. I hypothesize that another, perhaps more frequently used mechanism for island formation is from the segregation of existing genetic variation between populations experiencing different selection regimes. Herein I modeled genetic divergence from SGV to explore the parameter combinations likely to give rise to islands versus those that generate either genome-wide divergence or no divergence between populations. I considered seven different demographic scenarios that differ in terms of how SGV is partitioned within and between a pair of populations, the mating type, and the migration frequency between diverging populations. Our results highlight how the balance of migration and selection together with meta-population demography can strongly affect short term genome-wide patterns of differentiation.

## 2. Methods

### 2.1. Modeling divergence from standing genetic variation

I was interested in identifying the parameter range likely to give rise to islands (i.e., local differentiation only) from those that give rise to other genomic patterns (i.e., no or genome-wide differentiation). I considered scenarios in which 1) a pair of populations were completely isolated for a period of time that affected the partitioning of genetic variation between populations followed by 2) secondary contact and 3) divergence with gene flow. I was concerned here with SGV only and so I assumed that the genomic response to a given demographic scenario occurs without new mutations or that is on a shorter timescale than is relevant for new mutations. I examined the genome-wide and temporal dynamics of differentiation for 7 specific evolutionary scenarios that vary in how SGV is partitioned within and between populations, the degree of admixture between populations that occurred during secondary contact, the periodicity of migration, and whether individuals are obligate or facultative sexual (Table 1). In all scenarios, the initial type of SGV was a parameter of the model and I explore different levels of migration, divergent selection, and recombination.

The general life-history cycle during the divergence with gene flow following secondary contact is as follows. Migration between populations occurs at rate *m* between populations every *m_f_* generations. For obligate sexual cases, random mating occurs every generation, following migration if applicable. For facultative sexual cases, random mating occurs following migration only. In other words, for the facultative sexual scenarios, cell division occurs asexually and there are *m_f_* rounds of viability selection occurring between migration and random mating. Viability selection within populations occurs at the last step of the life cycle.

In each evolutionary scenario I tracked genetic differentiation between populations at neutral loci linked to a single locus under divergent selection. Locus *A* is under divergent selection between these two populations and it is linked to a neutral locus *B*. The dynamics of neutral divergence between populations can be tracked by following haplotype frequencies through time. Because there is only a single locus under selection, we can obtain genomic patterns of differentiation by varying the recombination rate, *r*, between loci *A* and *B*, migration between populations, and the strength of selection at locus *A*.

Let 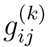 be the frequency of haplotypes in population *k* with allele *i* at selected locus *A* and allele *j* at neutral locus *B*. For convenience we can summarize the gamete frequencies for each population *k* as a vector, **p**_*k*_:

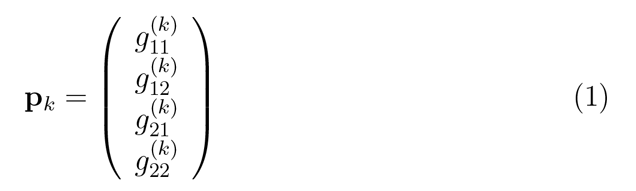

#### 2.1.1. Migration and random mating

Migration between the two populations experiencing divergent selection follows a simple two-island model with a migration rate *m*. For example, the vector of new haplotype frequencies following migration for population 1 is:

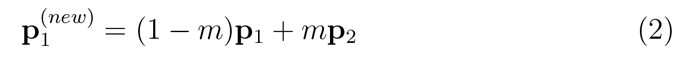

**Table 1:** Evolutionary models of explored in this study. For each scenario, the starting gamete frequencies, within-population LD, cross-population linkage disequilibrium (X-LD), mating mode, and frequency of migration (*m_f_*) is given. Admixture here refers to the number of random mating that occurred during the period of secondary contact.

| Scenario | mating | $m_f$ | Admixture | LD | X-LD | $g_{11}^{(1)}$ | $g_{12}^{(1)}$ | $g_{21}^{(1)}$ | $g_{22}^{(1)}$ | $g_{11}^{(2)}$ | $g_{12}^{(2)}$ | $g_{21}^{(2)}$ | $g_{22}^{(2)}$ |
| --- | --- | --- | --- | --- | --- | --- | --- | --- | --- | --- | --- | --- | --- |
| 1 | obligate | 1 | none | 0 | 0.25 | 1 | 0 | 0 | 0 | 0 | 0 | 0 | 1 |
| 2 | obligate | 1 | none | 0 | 0.125 | 0.5 | 0.5 | 0 | 0 | 0 | 0 | 0 | 1 |
| 3 | obligate | 1 | none | 0 | 0 | 0.5 | 0.5 | 0 | 0 | 0 | 0 | 0.5 | 0.5 |
| 4 | obligate | 1 | $F_1$s | 0.25 | 0.25 | 0.5 | 0 | 0 | 0.5 | 0.5 | 0 | 0 | 0.5 |
| 5 | obligate | 1 | $F_2$s | $0.25 - \frac{r}{4}$ | $0.25 - \frac{r}{4}$ | $0.5 - \frac{r}{4}$ | $\frac{r}{4}$ | $\frac{r}{4}$ | $0.5 - \frac{r}{4}$ | $0.5 - \frac{r}{4}$ | $\frac{r}{4}$ | $\frac{r}{4}$ | $0.5 - \frac{r}{4}$ |
| 6 | obligate | 50 | $F_2$s | $0.25 - \frac{r}{4}$ | $0.25 - \frac{r}{4}$ | $0.5 - \frac{r}{4}$ | $\frac{r}{4}$ | $\frac{r}{4}$ | $0.5 - \frac{r}{4}$ | $0.5 - \frac{r}{4}$ | $\frac{r}{4}$ | $\frac{r}{4}$ | $0.5 - \frac{r}{4}$ |
| 7 | facultative | 50 | $F_2$s | $0.25 - \frac{r}{4}$ | $0.25 - \frac{r}{4}$ | $0.5 - \frac{r}{4}$ | $\frac{r}{4}$ | $\frac{r}{4}$ | $0.5 - \frac{r}{4}$ | $0.5 - \frac{r}{4}$ | $\frac{r}{4}$ | $\frac{r}{4}$ | $0.5 - \frac{r}{4}$ |

Mating is assumed to occur at random amongst the individuals within a given population. The change in haplotype frequency after random mating is:

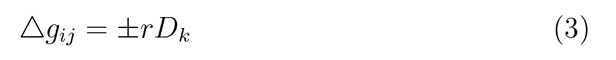
where *r* is the recombination rate between loci *A* and *B* and *D_k_* is the disequilibrium coefficient (*D_k_* = 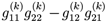). For the coupling gametes (i.e., *i* = *j*) the quantity *rD_k_* in Equation 3 is subtracted and it is added otherwise.

#### 2.1.2. Viability selection

A matrix describing the fitness values for all zygotes in population 1 is given by the matrix **S**_1_:

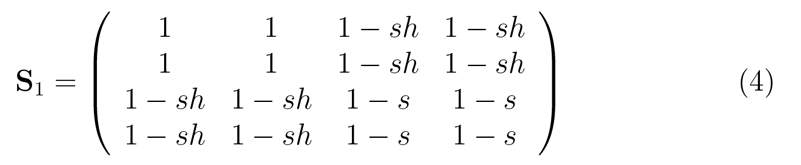
where *s* and *h* are the selection and dominance coefficients, respectively. For simplicity in the current model I assumed heterozygotes have intermediate fitness between the homozygote genotypes (i.e., *h* = 0.5). In equation 4 rows and columns correspond to the elements in **p**_*k*_. In population 2 the fitness matrix is constructed similarity but the quantity 1 – s is replaced with 1 and *vise versa*. The change in haplotype frequencies for each population can be calculated by considering the marginal fitness values for each haplotype. Following [12], the vector of marginal fitness values is:

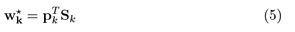

The change of haplotype frequencies due to viability selection depends on the mean relative fitness of a given population, the current haplotype frequency, and its marginal fitness. The mean relative fitness is the dot product of the haplotype frequencies and their corresponding marginal fitness values:

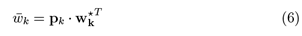

Thus, the vector of change of haplotype frequencies after a bout of selection is then:

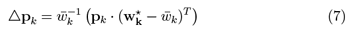

#### 2.1.3. Numerical methods

Since I was interested in the short-term dynamics of genomic differentiation following secondary contact, I ran each scenario for 500 generations for varying migration rates and strengths of selection and recorded the extent genetic differentiation [*F_ST_*, 4, 5] at each locus along a simulated chromosome.

## 3. Results

### 3.1. Genomic differentiation under DGF from secondary contact

#### 3.1.1. No admixture during secondary contact

Under the evolutionary scenarios in which no admixture during secondary contact occurred (i.e., scenarios 1-3, Table 1) I found that the extent of genetic differentiation depended on the relative magnitudes of migration and selection (Figure 1). When initial divergence was strong (i.e., scenario #1) the small increases in the migration rate greatly reduced overall differentiation in about 100 generations. Here, under weak to intermediate migration (i.e., 0.001 ≥ *m* ≥ 0.01) and under intermediate to strong selection (i.e., *s* ≥ 0.05) genomic differentiation occurred only under tight linkage, consistent with genomic islands. This same general pattern was observed when the initial divergence was weaker (LD=0, X-LD=0.125, scenario #2, Figure 2) but with less overall differentiation. As expected, when SGV was partitioned completely within populations no differentiation occurred in any migration and selection range (scenario #3, Figure 3).

#### 3.1.2. Brief admixture during secondary contact

I found that brief admixture between diverged, locally adapted populations immediately before DGF strongly promoted island formation. Indeed, when DGF was initiated with *F*_1_ individuals – the parents of which were locally adapted to their respective environment – I found that the only divergence that was detected occurred locally within the genome (scenario #4, Figure 4). This island pattern was also observed when two rounds of random mating occurred prior to DGF (scenario #5, Figure 5).

I found a strong effect of mating type on the pattern of genetic differentiation from SGV. As predicted, for obligate sexual mating and when migration occurs periodically (e.g., every 50 generations, scenario #6) selection is relatively strong compared to migration resulting in island formation and persistence even under maximum migration (m = 0.5; migration per generation = 0.01). When mating type is facultative, however, the joint contribution of selection and migration can create genome-wide differentiation in addition to islands (Figure 7). Here, genome-wide differentiation occurs under strong selection and weak migration.

I identified two pathways in which islands form, depending on the relative strength of selection and migration. First, under strong selection (*s* ≥ 0.05) and strong migration (*m* ≥ 0.2) islands form from the breakdown of genomic differentiation with time (e.g., upper right panels of Figures 6 and 7). Second, under weak selection (s=0.01) and weak to moderate migration (0.01 < *m* < 0.05), neutral genetic differentiation began low and increased (“grew”) over time (e.g., Figures 6 and 7). The size (width) of islands differed between the two mating types – with larger islands found in facultative compared to obligate mating types. Interestingly, migration was not need for island growth to occur when admixture occurred during a single bout of secondary contact (*m* = 0, *s* = 0.01, Figures 4, 5, 6, and 6). This is in stark contrast to scenarios in which no admixture occurred in secondary contact (scenarios 1-3, Figures 1, 2, and 1).

## 4. Discussion

### 4.1. Islands from standing genetic variation

I found that localized genetic differentiation can readily occur under a wide range of demographic scenarios, depending on the relative strength of migration and divergent selection. Linkage disequilibrium within and between isolated populations can be generated a number of ways prior to the onset of a divergent selection regime. For example, genetic drift can fix alternative alleles between two isolated populations such that there is no LD within but maximum LD between populations. Of course, the fixation of alternative alleles in each isolated population can occur due to preexisting divergent selection on new mutations. In general, the breakdown of linkage disequilibrium under divergence is required for islands to form.

### 4.2. Islands uplift and islands erode

Under new mutation theory of island formation, divergent hitchhiking allows for increase establishment probability of new mutations [17] and so islands can “uplift” from the metaphorical sea when seeded with divergently selected loci. I found that such uplifting can also occur from standing genetic variation. An admixture event between genotypically distinct populations creates a high degree of within population LD [5]. Such a case may occur between hybridizing sister species or through the ephemeral breakdown of a migration barrier. When divergent selection occurs following such an event there are two mechanisms in which islands can form, depending the strength of selection relative to gene flow. During the time in which LD is broken down within a population by random mating, differentiation at both selected and neutral loci increases (though this increase is faster at the selected locus; Figure 8E-F). In the case of no migration between populations (e.g., left column of Figure 8), neutral differentiation will remain steady since no migration (or mutation) is occurring. Islands can also buildup quickly and erode. For example, under strong divergence with moderate gene flow there is a rapid breakdown of LD early with a slower breakdown of LD later (Figure 8D). During the rapid breakdown phase, where the change in haplotype frequencies is dominated by selection and *F_ST_* increases with time for both selected and linked neutral sites. During the slow breakdown of LD phase, the change in haplotype frequencies are dominated by migration. Here, differentiation at the selected locus is stable whereas differentiation decreases at the neutral locus (Figure 8F) owing to recombination. With tighter (weaker) linkage the decrease of *F_ST_* will be slower (higher). Thus, under divergence with gene flow I would expect islands to erode with time.

**Figure 1:**
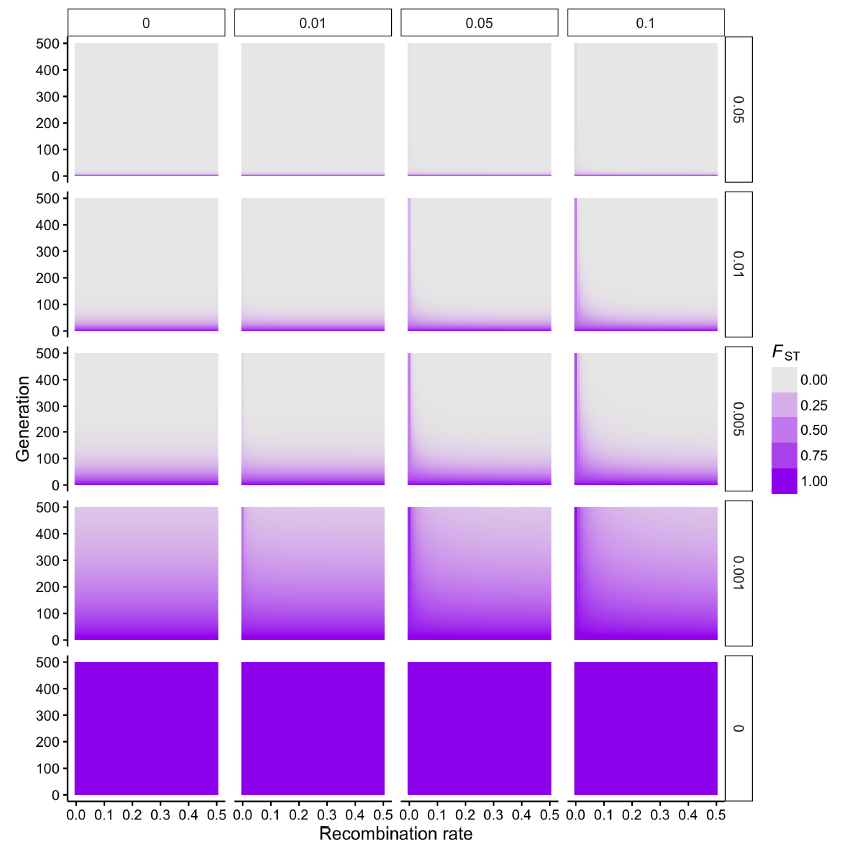
Dynamics of divergence with gene flow under scenario #1 – obligate sexual, *m_f_* = 1, LD=0, X-LD=0.25. For each panel, the extent of divergence (*F_ST_*) at neutral loci are given across time. Rows indicate migration rate, *m*, between diverging populations and columns indicate the strength of divergent selection, *s*, at selected locus *A*.

**Figure 2:**
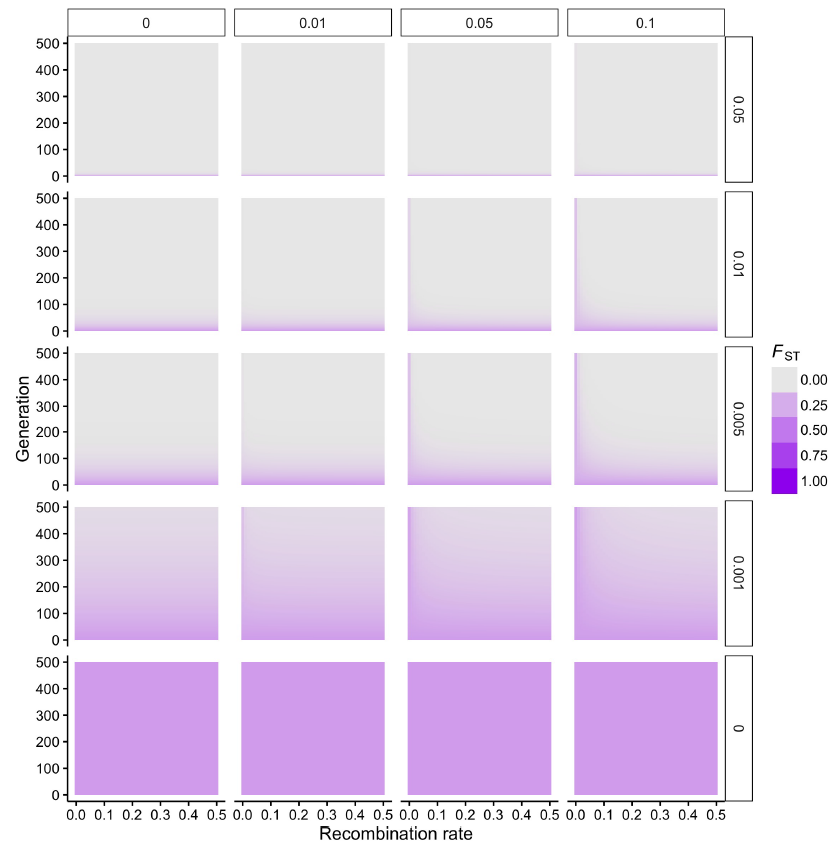
Dynamics of divergence with gene flow under scenario #2 – obligate sexual, *m_f_* = 1, LD=0, X-LD=0.125. For each panel, the extent of divergence (*F_st_*) at neutral loci are given across time. Rows indicate migration rate, *m*, between diverging populations and columns indicate the strength of divergent selection, *s*, at selected locus *A*.

**Figure 3:**
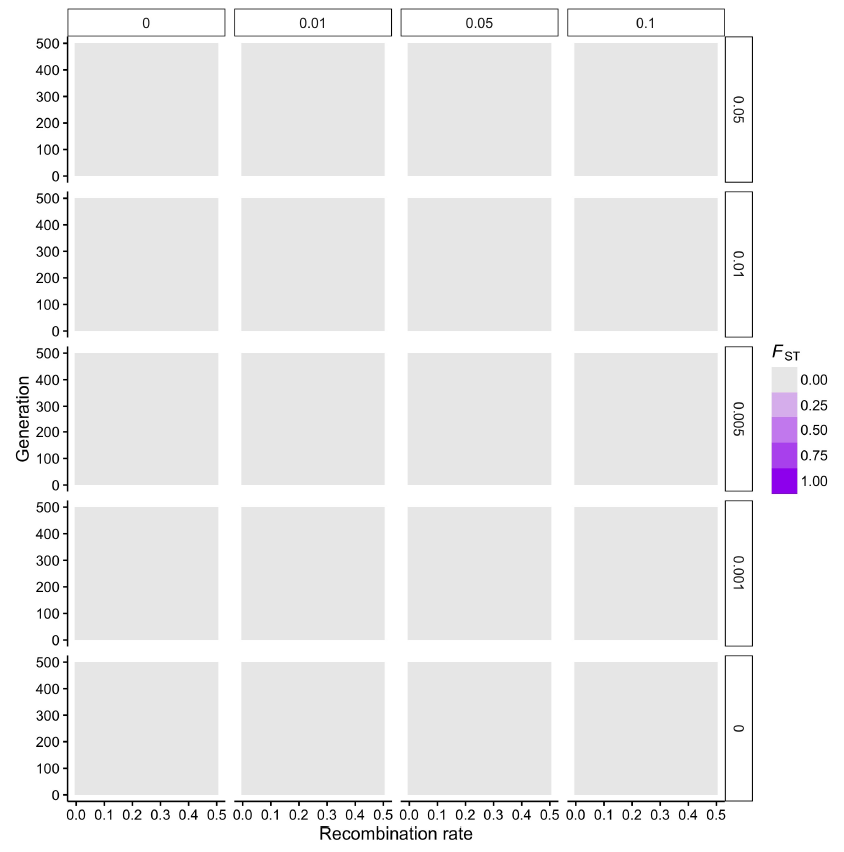
Dynamics of divergence with gene flow under scenario #3 – obligate sexual, *m_f_* = 1, LD=0, X-LD=0. For each panel, the extent of divergence (*F_ST_*) at neutral loci are given across time. Rows indicate migration rate, *m*, between diverging populations and columns indicate the strength of divergent selection, *s*, at selected locus *A*.

**Figure 4:**
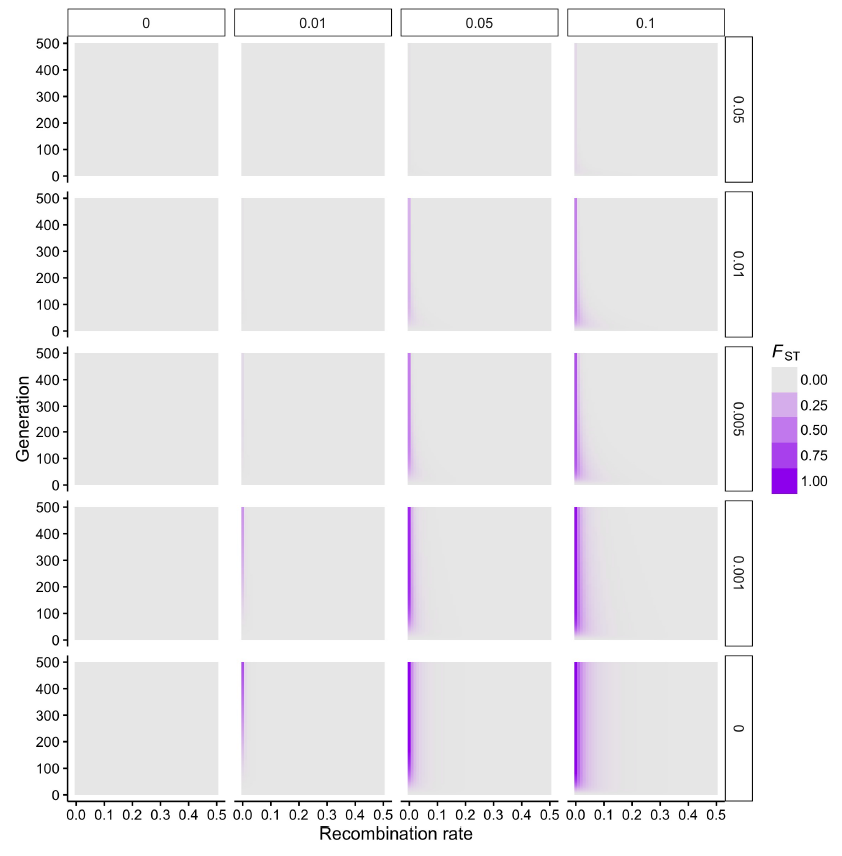
Dynamics of divergence with gene flow under scenario #4 – obligate sexual, *m_f_* = 1, LD=0.25, X-LD=0.25. For each panel, the extent of divergence (*F_ST_*) at neutral loci are given across time. Rows indicate migration rate, *m*, between diverging populations and columns indicate the strength of divergent selection, *s*, at selected locus *A*.

**Figure 5:**
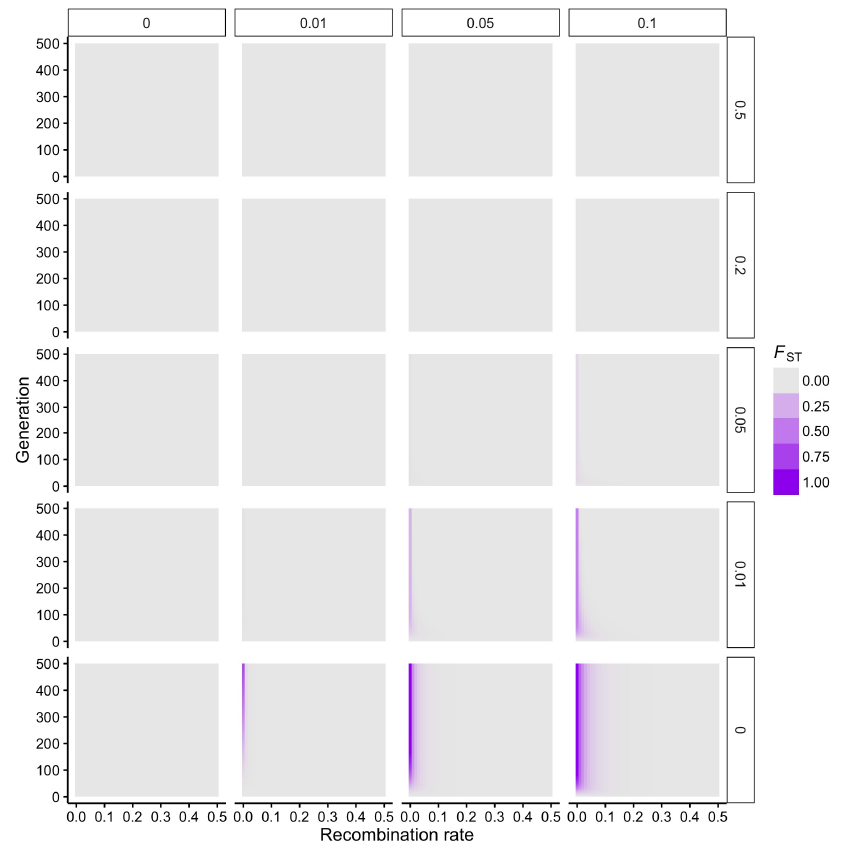
Dynamics of divergence with gene flow under scenario #5 – obligate sexual, *m_f_* = 1, LD=0.25 - 0.25r, X-LD=0.25 - 0.25r. For each panel, the extent of divergence (*F_st_*) at neutral loci are given across time. Rows indicate migration rate, *m*, between diverging populations and columns indicate the strength of divergent selection, *s*, at selected locus *A*. Note the change of migration rates investigated in this figure compared to Figure 4

**Figure 6:**
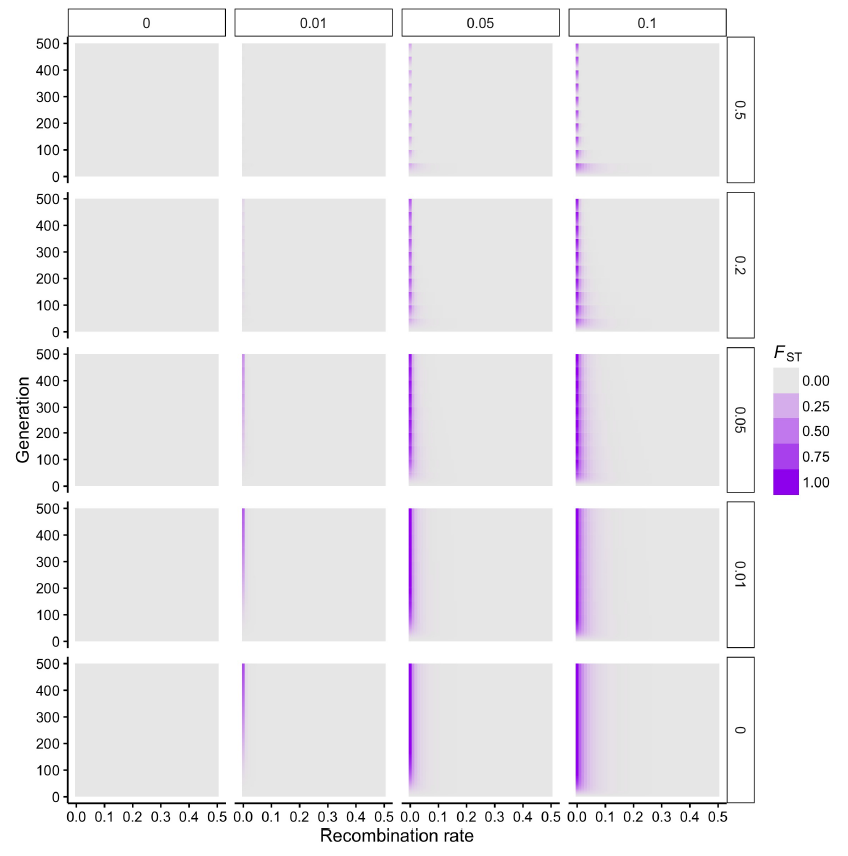
Dynamics of divergence with gene flow under scenario #6 – obligate sexual, *m_f_* = 1, LD=0.25 - 0.25r, X-LD=0.25 - 0.25r. For each panel, the extent of divergence (*F_st_*) at neutral loci are given across time. Rows indicate migration rate, *m*, between diverging populations and columns indicate the strength of divergent selection, *s*, at selected locus *A*.

**Figure 7:**
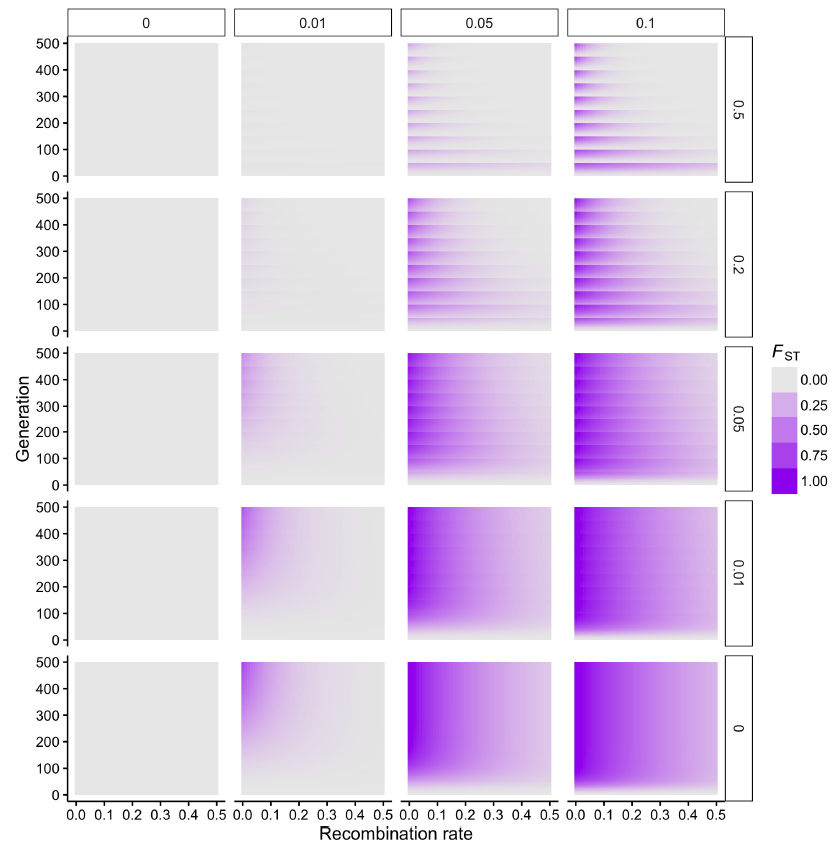
Dynamics of divergence with gene flow under scenario #7 – facultative sexual, *m_f_* = 50, LD=0.25 - 0.25r, X-LD=0.25 - 0.25r. For each panel, the extent of divergence (*F_ST_*) at neutral loci are given across time. Rows indicate migration rate, *m*, between diverging populations and columns indicate the strength of divergent selection, *s*, at selected locus *A*.

**Figure 8:**
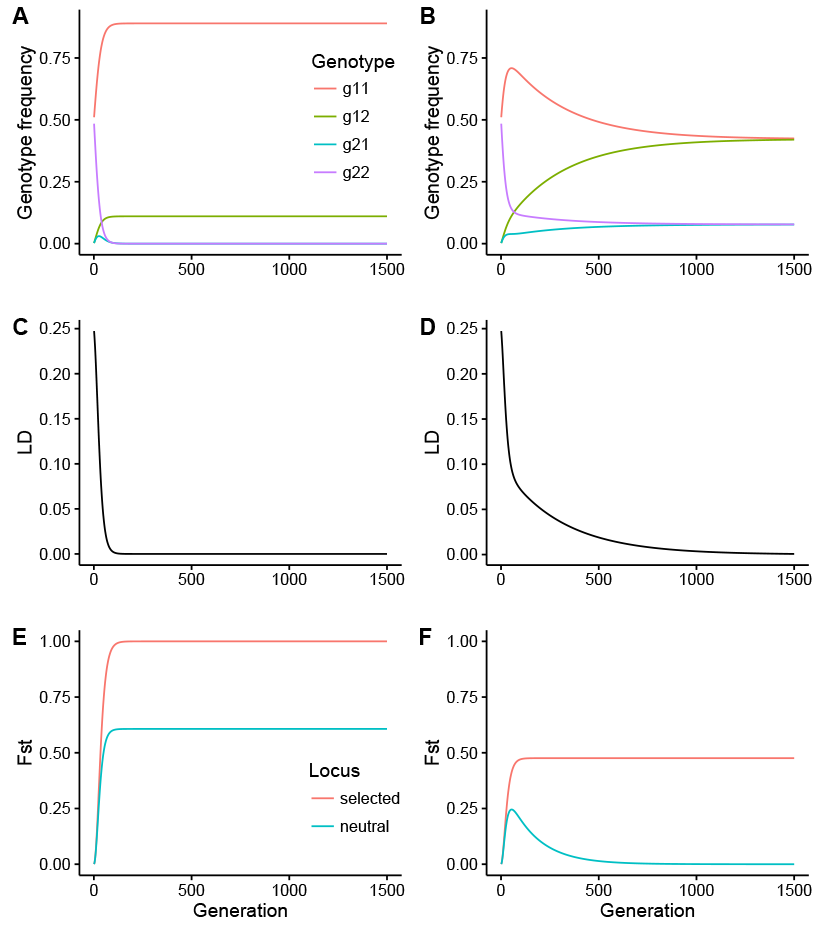
Temporal dynamics of genotype frequencies, LD, and differentiation at the selected and linked neutral loci (scenario #4). Two specific examples (left and right columns) of island formation are given. For each example I plotted the results for population #1 only so that genotypes *g*_11_ and *g*_12_ are favored and *g*_22_ and *g*_21_ are disfavored. Left column, migration is absent. Right column, migration is weak (*m* = 0.01). For each condition the selection coefficient was strong (*s* = 0.1) and the recombination rate between the selected and neutral loci was 0.01.

